# National Institutes of Health Research Project Grant Inflation 1998 to 2021

**DOI:** 10.1101/2022.10.07.511257

**Authors:** Michael Lauer, Joy Wang, Deepshikha Roychowdhury

## Abstract

We analyzed changes in total costs for National Institutes of Health (NIH) awarded Research Project Grants issued from 1998 to 2003. The NIH uses a data-driven price index – the Biomedical Research and Development Price Index (BRDPI) – to account for inflation based increases in grant costs. The BRDPI was higher than the general rate of inflation from 1998 until 2012; since then, the rate of inflation for NIH-funded research has been similar to the general rate of inflation. Despite increases in nominal costs, recent years have seen increases in the absolute numbers of RPG and R01 awards. Real (BRDPI-adjusted) average and median RPG costs increased during the NIH-doubling (1998 to 2003), but have remained relatively stable since. Of note, though, the degree of variation of RPG costs has changed over time, with more marked extremes observed on both higher and lower levels of cost. On both ends of the cost spectrum, the agency is funding a greater proportion of solicited projects, with nearly half of RPG money going towards solicited projects. After adjusting for potential confounders in a wholly non-parametric machine learning regression, we find no independent association of time with BRDPI-adjusted costs.

## Introduction

Inflation, defined by the United States Federal Reserve as “the increase in the prices of goods and services over time”(US Federal Reserve, 2022), has been a longstanding concern in the biomedical research community.(Mervis, 2015) Concern has only increased over the past year given the increased rate of inflation in the general economy.

The National Institutes of Health (NIH) issues different type of research and training awards, but by far the most common type is the “Research Project Grant (RPG)”(NIH, 2022a) accounting for over half of the NIH budget.(NIH, 2022b) Prices for research project grants (RPGs) awarded by the National Institutes of Health (NIH) may increase over time for at least three reasons:

- Background inflation: Increases in prices across the economy due to increases in the money supply and/or economy-wide demand and supply stresses; these are reflected in general price indices, such as the GDP price index(NIH, 2022c) and the Consumer Price Index.(US Bureau of Labor Statistics, 2022)
- Research-specific inflation: Increases in prices in the biomedical research and development enterprise; these are reported as the Biomedical Research and Development Price Index (or BRDPI).(NIH, 2022c) The BRDPI measures changes in the weighted average of the prices of all the inputs (e.g., personnel services, various supplies, and equipment) purchased with the NIH budget to support research. The weights used to construct the index reflect the actual pattern, or proportions,of total NIH expenditures on each of the types of inputs purchased. Theoretically,the annual change in the BRDPI indicates how much NIH expenditures would need to increase, without regard to efficiency gains or changes in government priorities,to maintain NIH-funded research activity at the previous year’s level.
- Changes in agency purchasing decisions (or compositional effects): We might imagine an automobile-rental firm that starts one year purchasing 10 mid-size sedans. The following year, it might choose to purchase instead 10 luxury mid-size sedans; costs increase not because of background inflation because of the firm’s decisions about what it wants to buy. Alternatively, the firm may purchase 2 large vans, 4 mid-sized sedans, and 4 compact cars. Overall and median costs might not change (compared to the baseline of 10 mid-size sedans), but the firm’s management will be acutely aware of the costs of the 2 large vans. Similarly, NIH Institutes and Centers (IC’s) may choose to “puchase”investigator-initiated R01 awards, R01 awards that cost more (e.g. >500K in direct costs) because of use of large animals, or different size awards (program project grants, cooperative agreements, or small exploratory R21 or R03 awards).

We report on the distribution of nominal and inflation-adjusted prices of funded NIH RPGs since 1998, the year that the NIH budget doubling began. We find that median and mean inflation-adjusted RPG costs have been largely stable since the doubling ended in 2003, but that there have been changes in the distribution (variance) of costs, which largely reflect compositional effects as agency priorities have shifted over time.

## Results

### Changes in RPG Costs and Characteristics over Time

Most of this report will focus on real (as opposed to nominal) costs of NIH RPG awards, that is total costs per RPG indexed for the FY2021 BRDPI. For context Figure 1, panel A, shows the number of unique parent competing and non-competing RPGs awarded each year. Between 1998 and 2021, NIH issued 827,815 RPG awards of at least $25,000 per year (BRDPI-indexed to 2021). The 3 vertical dotted lines refer to the beginning and end of the NIH-doubling (1998 and 2003) and the year of budget sequestration (2013). The number of RPGs increased during the doubling, decreased gradually between the end of the doubling and 2015, and increased again with recent NIH budget increases. Figure 1, panel B, shows the GDP Price Index and the BRDPI. Between 1998 and 2012 the BRDPI was consistently higher than the GDP Price Index; after 2012, when the government imposed caps on compensation of extramural investigators, the BRDPI has fallen to the same levels as the GDP Price Index.

**Figure 1:**
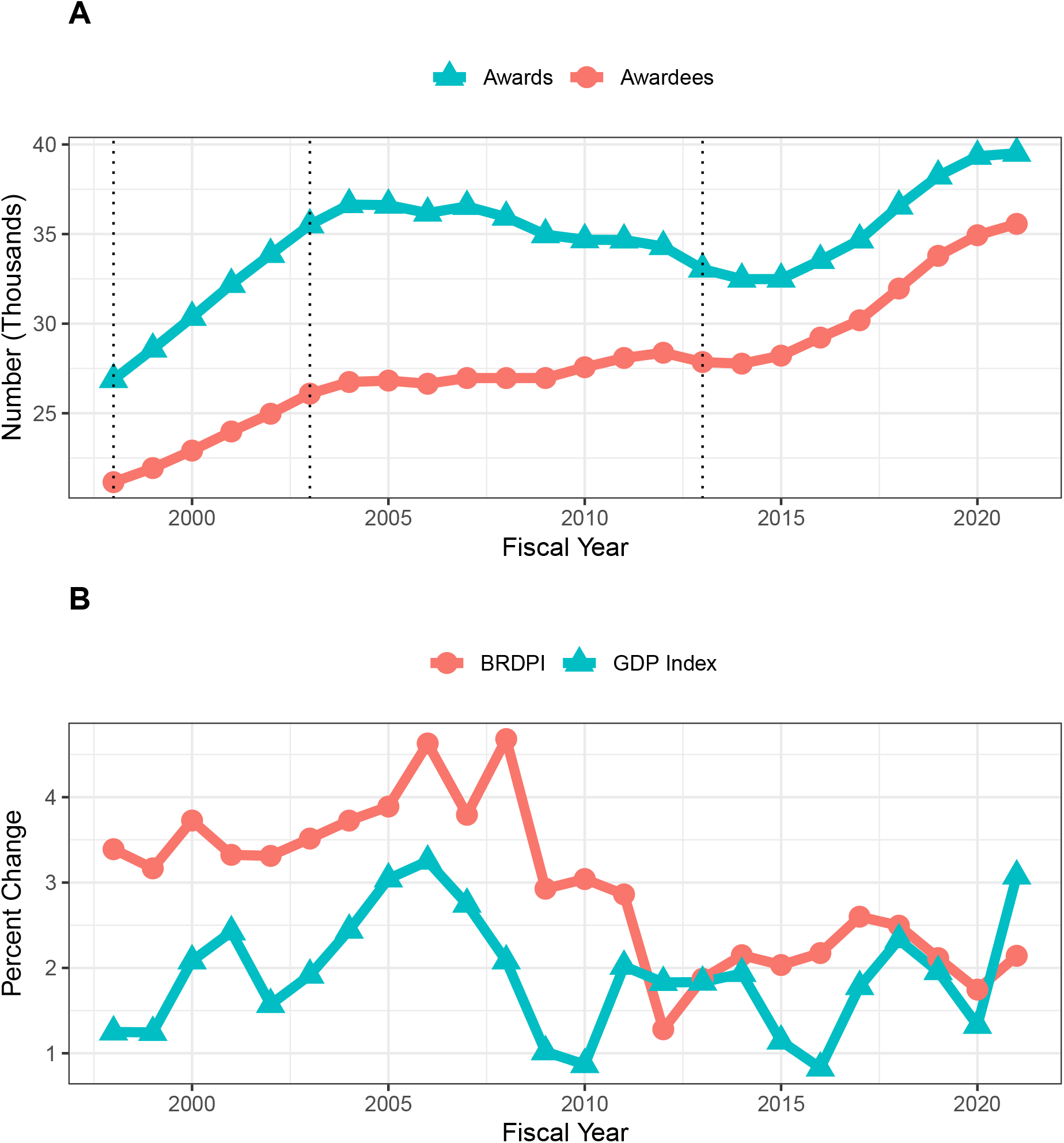
Number of funded RPGs and RPG awardees (panel A) and inflationary indices (panel B) by fiscal year

### Nominal and BRDPI-Indexed Costs of NIH RPGs over Time

Figure 2 shows mean (panel A) and median (panel B) costs per RPG in both nominal (red lines with circles) and FY2021 BRDPI-adjusted (teal lines with triangles) values. The real (that is FY2021 BRDPI-adjusted) costs of RPGs increased during the doubling (from average values of about $530,000 to about $610,000), fell to a nadir of about $520,000 in 2013, and after a quick rebound in 2014 has remained relatively stable at about $570,000 since. Figure 3 focuses on R01 awards and shows that the number of R01 awards has increased over recent years to the levels seen with the NIH doubling. Figure 4 shows mean (panel A) and median (panel B) costs per R01 award in nominal (red lines with circles) and FY2021 BRDPI-adjusted (teal lines with triangles) values. Trends are similar as with RPG awards overall (Figure 2).

**Figure 2:**
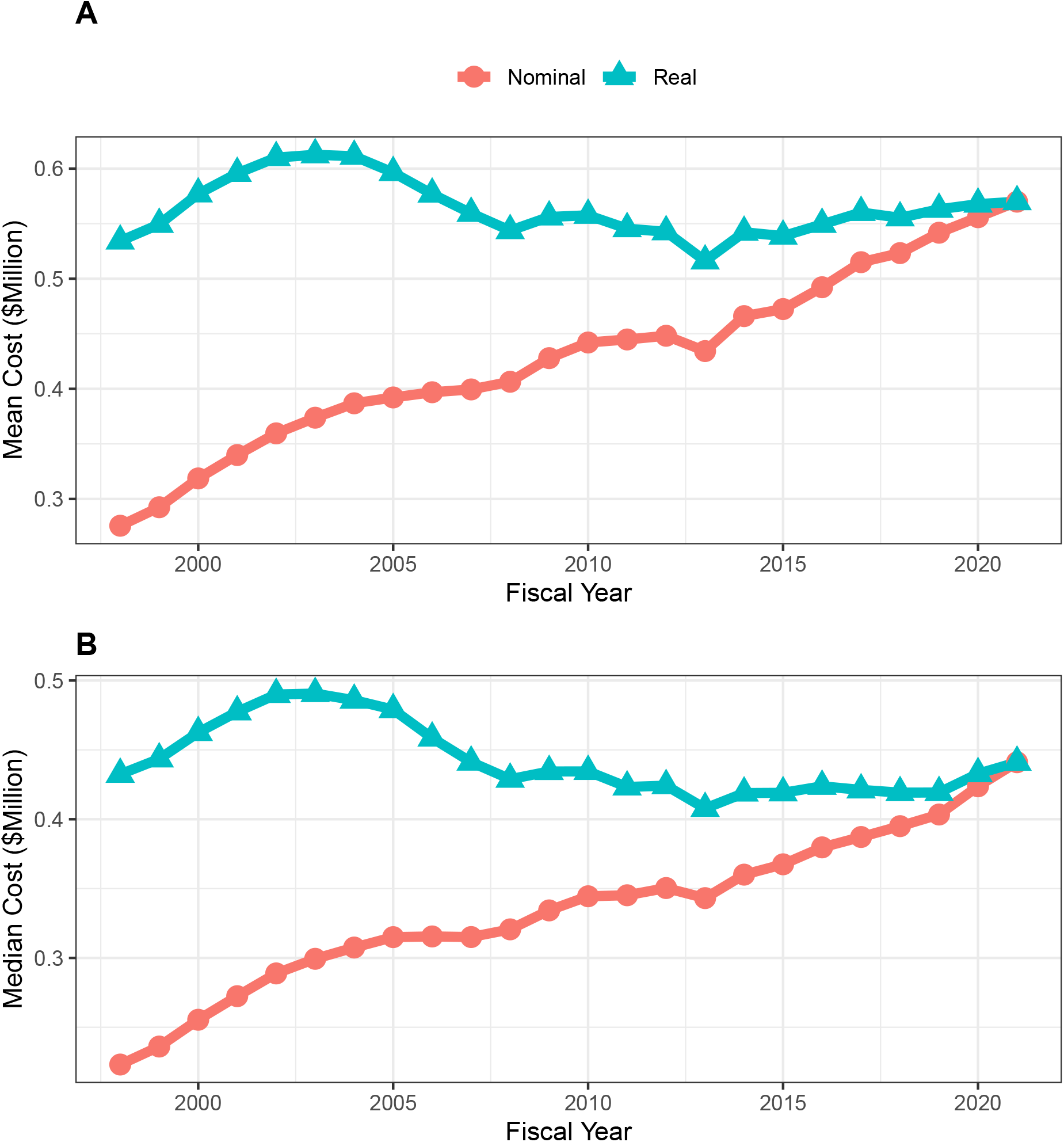
Mean (panel A) and median (panel B) nominal and real (BRDPI-adjusted) costs for NIH-funded RPGs, FY1998 to FY2021.

**Figure 3:**
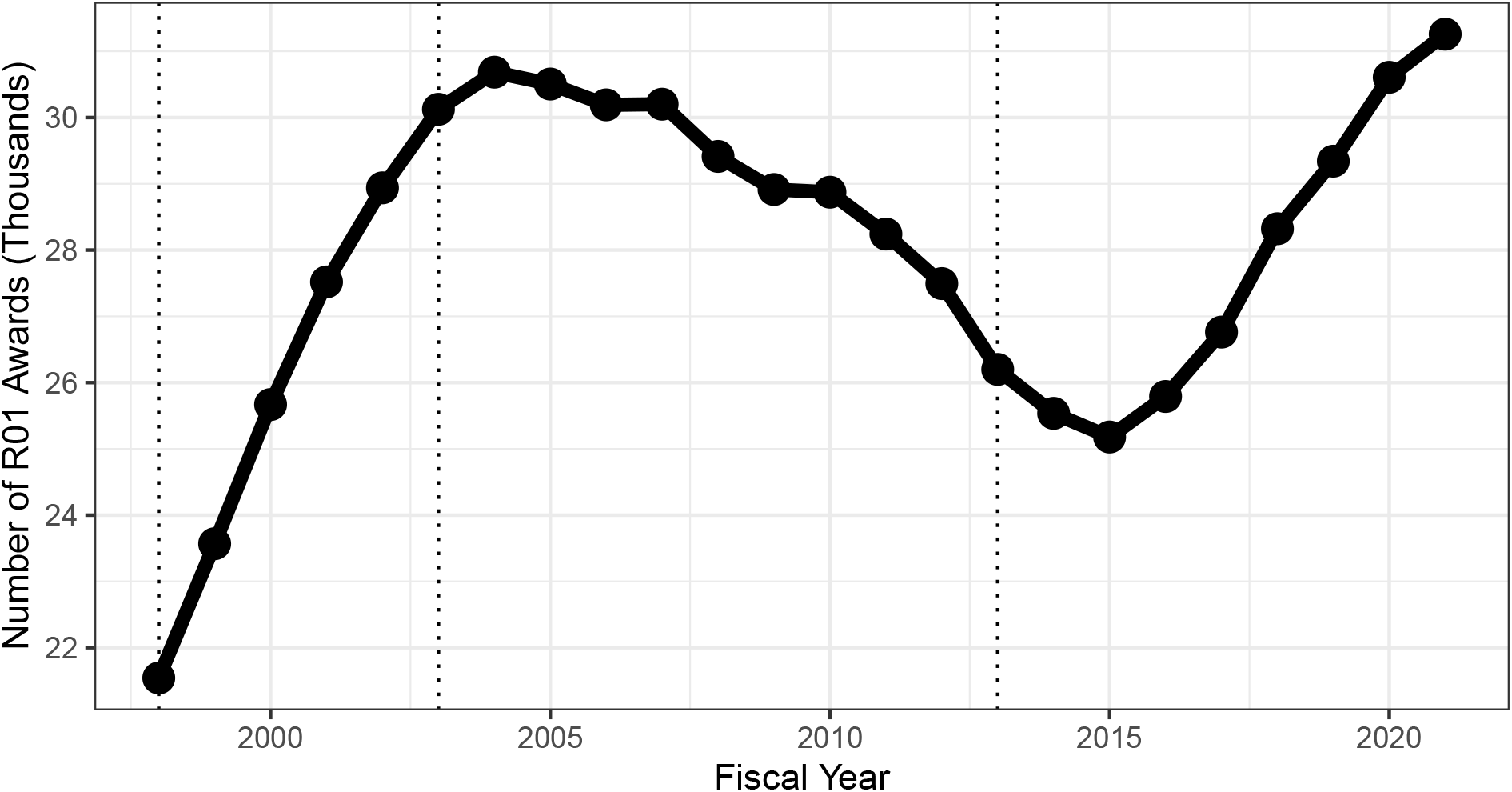
Number of funded R01-equivalent awards by fiscal year

**Figure 4:**
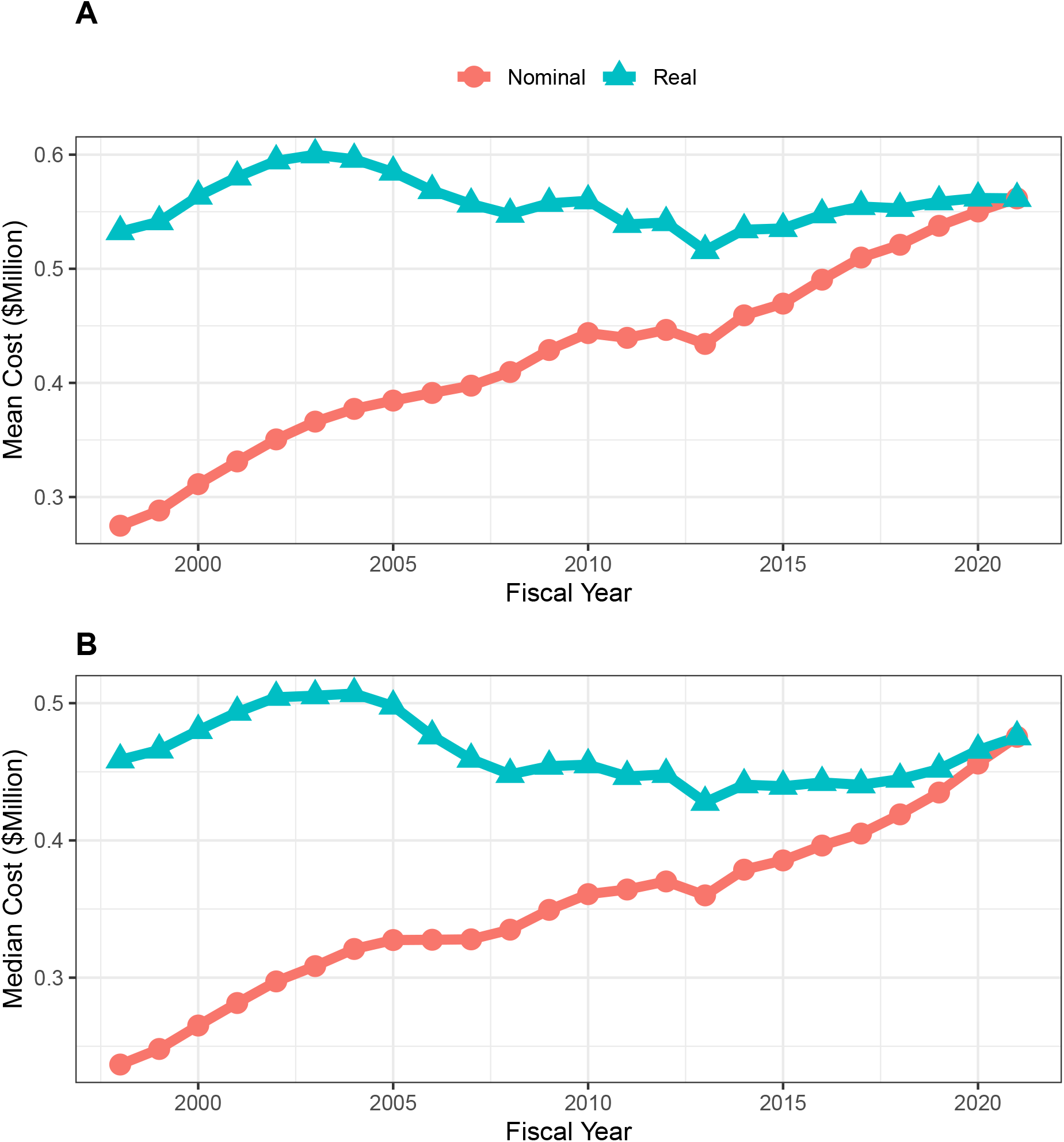
Mean (panel A) and median (panel B) nominal and real (BRDPI-adjusted) costs for NIH-funded R01-equivalent awards, FY1998 to FY2021.

### Characteristics of NIH RPGs over Time

Table 1 shows characteristics of NIH RPGs in selected years. Over time there have been decreases in the proportion of unsolicited awards and program (“P”) grants, while there have been increases in the proportions of R21 or R03 grants, cooperative agreements, clinical trials (since reliable data were first collected in 2008), highly expensive projects (defined as those costing at least $5 million in FY2021 BRDPI-adjusted, *not nomimal*, values), and human-studies only projects. The proportion of R01-equivalent awards increased during the doubling and then returned to 1998 levels. Institutes of higher education, independent research organizations, and independent hospitals have consistently accounted for over 97 percent of awards.

**Table 1:**
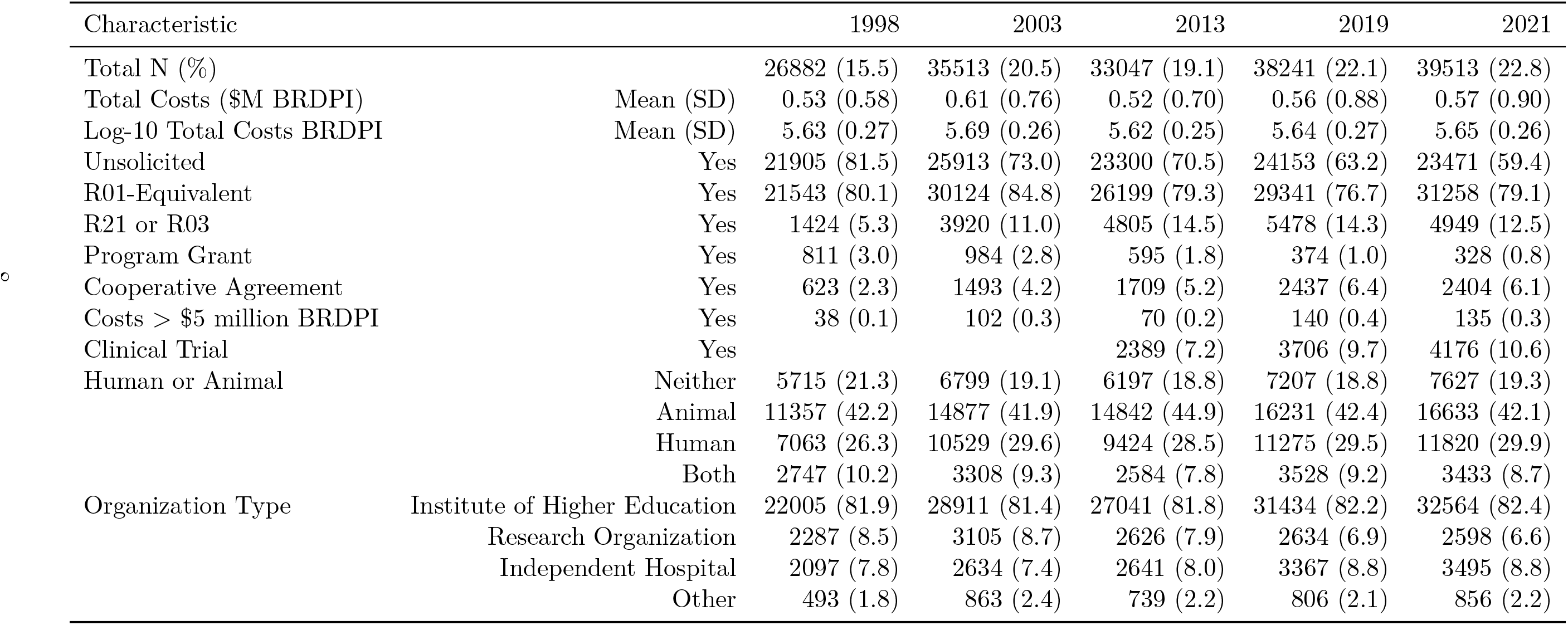
Characteristics of RPGs in selected fiscal years.

| Characteristic |  | 1998 | 2003 | 2013 | 2019 | 2021 |
| --- | --- | --- | --- | --- | --- | --- |
| Total N (%) |  | 26882 (15.5) | 35513 (20.5) | 33047 (19.1) | 38241 (22.1) | 39513 (22.8) |
| Total Costs (\$M BRDPI) | Mean (SD) | 0.53 (0.58) | 0.61 (0.76) | 0.52 (0.70) | 0.56 (0.88) | 0.57 (0.90) |
| Log-10 Total Costs BRDPI | Mean (SD) | 5.63 (0.27) | 5.69 (0.26) | 5.62 (0.25) | 5.64 (0.27) | 5.65 (0.26) |
| Unsolicited | Yes | 21905 (81.5) | 25913 (73.0) | 23300 (70.5) | 24153 (63.2) | 23471 (59.4) |
| R01-Equivalent | Yes | 21543 (80.1) | 30124 (84.8) | 26199 (79.3) | 29341 (76.7) | 31258 (79.1) |
| R21 or R03 | Yes | 1424 (5.3) | 3920 (11.0) | 4805 (14.5) | 5478 (14.3) | 4949 (12.5) |
| Program Grant | Yes | 811 (3.0) | 984 (2.8) | 595 (1.8) | 374 (1.0) | 328 (0.8) |
| Cooperative Agreement | Yes | 623 (2.3) | 1493 (4.2) | 1709 (5.2) | 2437 (6.4) | 2404 (6.1) |
| Costs > \$5 million BRDPI | Yes | 38 (0.1) | 102 (0.3) | 70 (0.2) | 140 (0.4) | 135 (0.3) |
| Clinical Trial | Yes |  |  | 2389 (7.2) | 3706 (9.7) | 4176 (10.6) |
| Human or Animal | Neither | 5715 (21.3) | 6799 (19.1) | 6197 (18.8) | 7207 (18.8) | 7627 (19.3) |
|  | Animal | 11357 (42.2) | 14877 (41.9) | 14842 (44.9) | 16231 (42.4) | 16633 (42.1) |
|  | Human | 7063 (26.3) | 10529 (29.6) | 9424 (28.5) | 11275 (29.5) | 11820 (29.9) |
|  | Both | 2747 (10.2) | 3308 (9.3) | 2584 (7.8) | 3528 (9.2) | 3433 (8.7) |
| Organization Type | Institute of Higher Education | 22005 (81.9) | 28911 (81.4) | 27041 (81.8) | 31434 (82.2) | 32564 (82.4) |
|  | Research Organization | 2287 (8.5) | 3105 (8.7) | 2626 (7.9) | 2634 (6.9) | 2598 (6.6) |
|  | Independent Hospital | 2097 (7.8) | 2634 (7.4) | 2641 (8.0) | 3367 (8.8) | 3495 (8.8) |
|  | Other | 493 (1.8) | 863 (2.4) | 739 (2.2) | 806 (2.1) | 856 (2.2) |

### Variation in RPG Costs over Time

Figure 5, panel A, shows box-plot distributions of FY2021 BRDPI-adjusted total cost per RPG over time. The means (shown as diamonds) and medians correspond to the data shown in Figure 2, panels A (means) and B (medians). Of note, the means are much greater than the medians, consistent with highly skewed distributions. The whiskers are also quite long, consistent with fat-tailed distributions. We can address skewness by log transforming BRDPI-adjusted total costs (*TC*), that is calculating *log_10_TC_BRDPI_*, with results shown in Figure 5, panel B. With log-transformation means and medians are nearly equal (eliminating skewness), but the whiskers remain prominent reflective of fat tails on both more expensive and less expensive ends.

**Figure 5:**
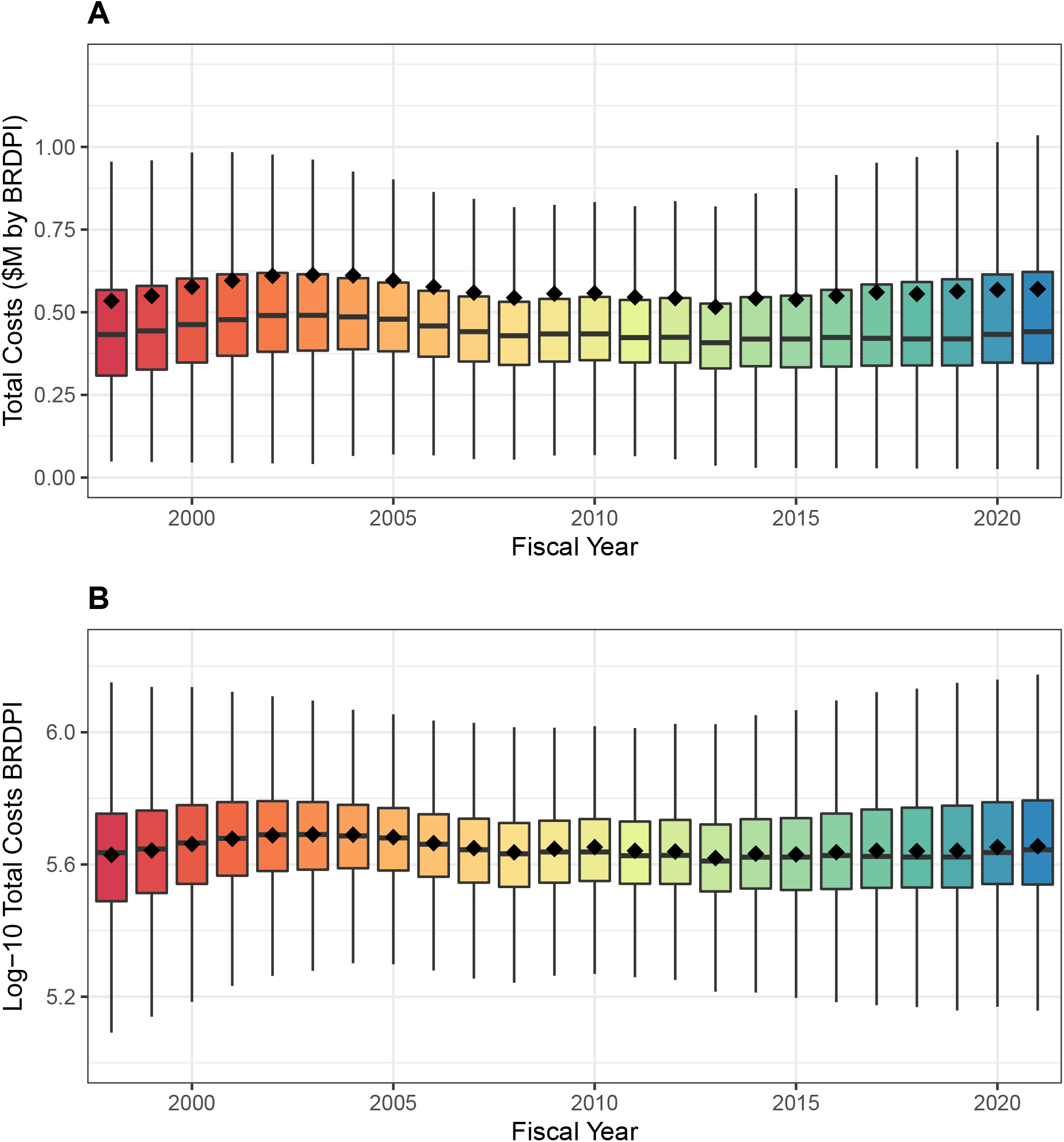
Box plot distributions of total costs (panel A) and log-transformed costs (panel B) for NIH-funded RPGs, FY1998 to FY2021. The diamonds refer to mean values. All costs are FY2021 BRDPI-adjusted.

Careful inspection of both panels in Figure 5 reveals an interesting pattern in variation. From the time of the doubling until about 2010, the distance between the whisker tips decreased. We can call this distance the “whisker range.” From 2012 through 2021 whisker ranges increased, exceeding levels for the doubling for untransformed costs, and not quite reaching doubling levels for log-transformed costs. We can think of the upper (and lower) whisker tips as the most extremely expensive (inexpensive) award that is not an outlier; the distance of the tips from the center (median) reflects the agency’s general willingness to vary its funding instruments. Figure 6 shows the whisker ranges declined from $920,000 to $750,000 between FY2002 and FY2010 and increased to over $1 million in FY2021 (panel A, with log-transformed values shown in panel B).

**Figure 6:**
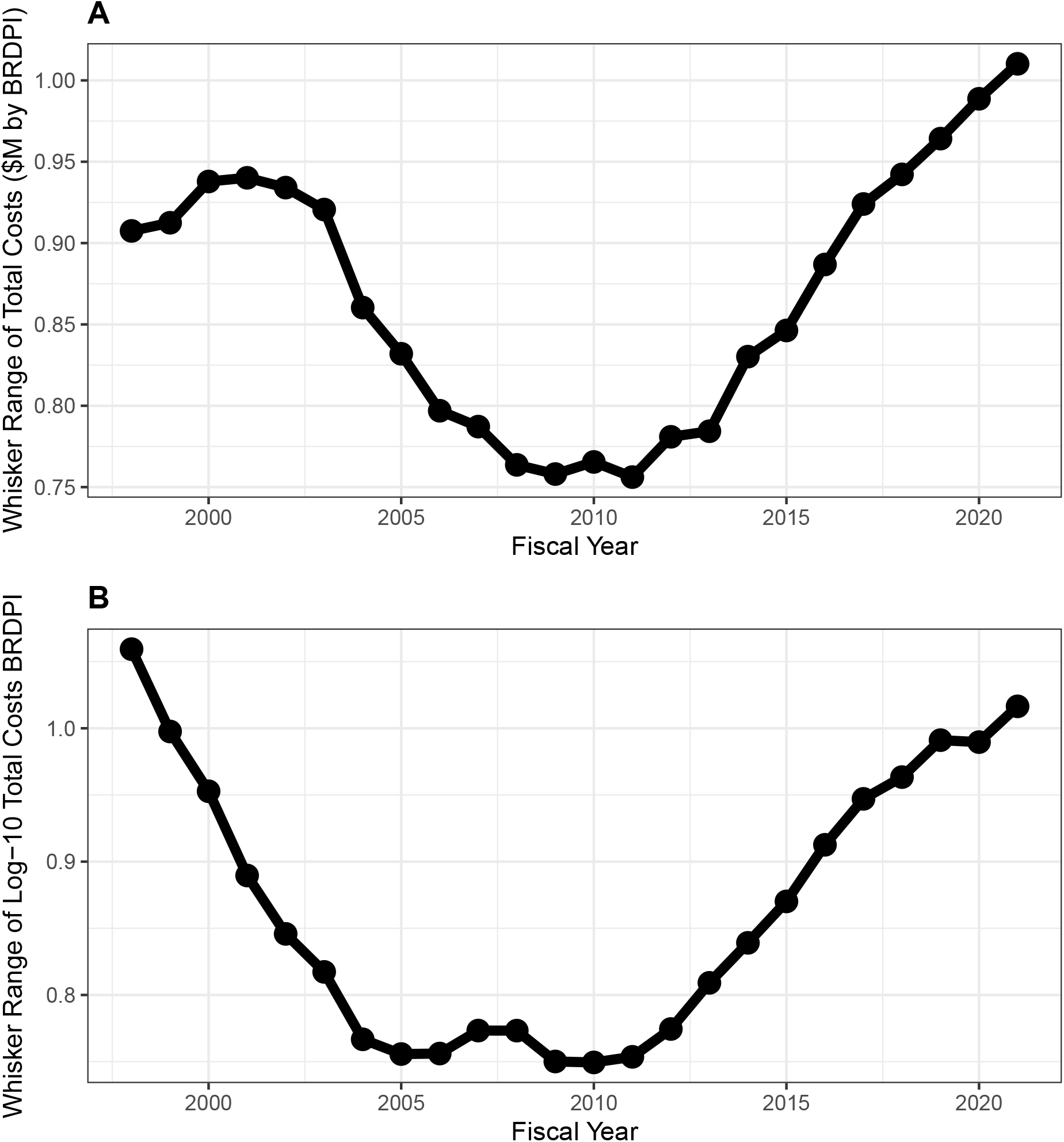
Whisker ranges, that is the distance between the tips of upper and lower whiskers, of box plots shown in Figure 5. All costs are FY2021 BRDPI-adjusted.

What might be behind the increasing extremes (higher and lower) over the past 10−15 years? Figure 7 shows the proportion of all RPG funding going to the top (bottom) centile and decile of costs. In 1998, the top centile (panel A) of RPG awards received 8 percent of funding, rising to 12 percent 2017; this 4 percent absolute difference means that an additional $850 million were awarded to approximately 1650 grants. There was little change in the proportion of funding going to the top decile; thus the upper extreme seems to be driven by increases in funding going to the most expensive awards.

**Figure 7:**
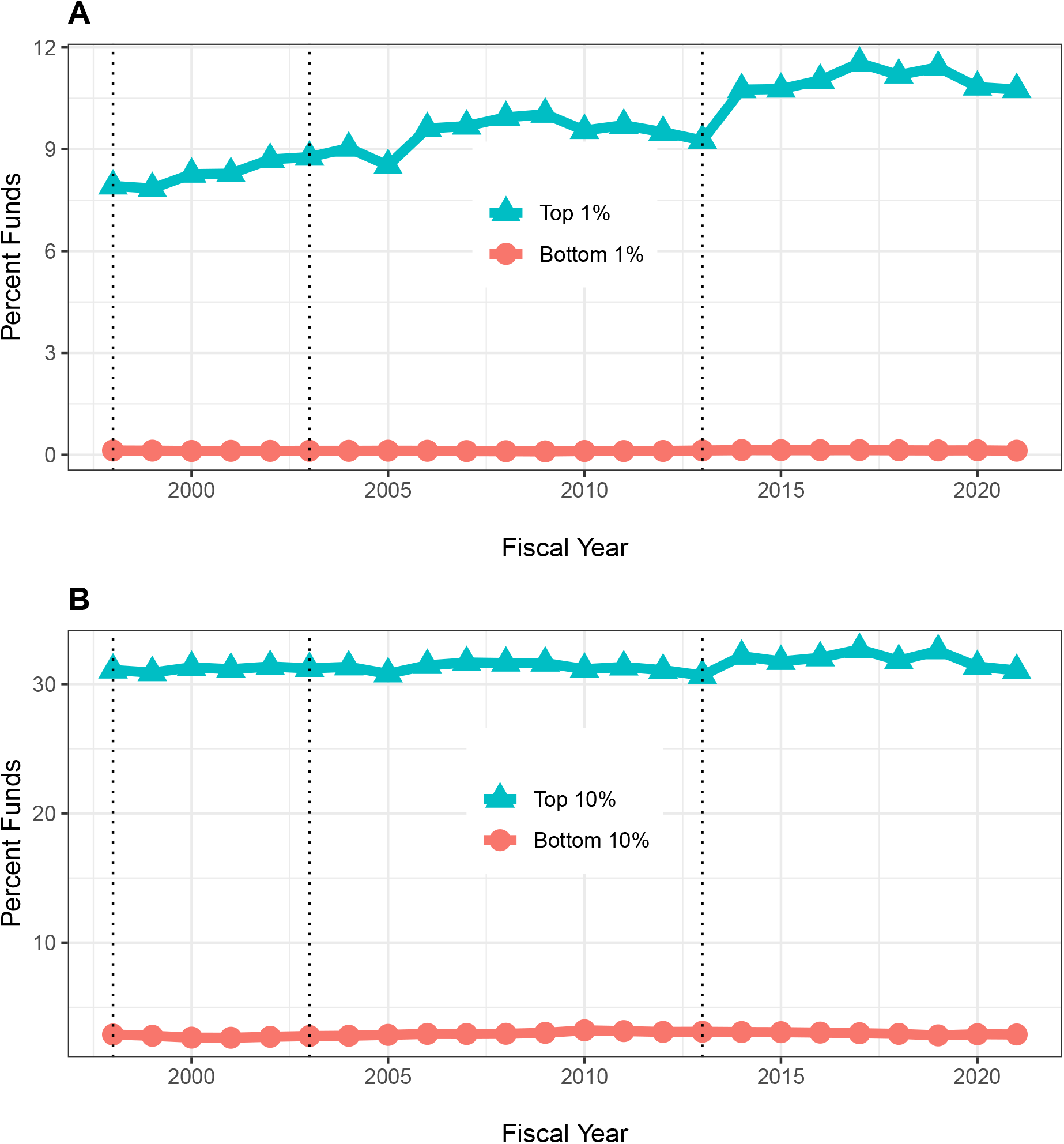
Distribution of total funds to the top and bottom centiles (panel A) and deciles (panel B) of NIH-funded RPGs, FY1998 to FY2021.

### Solicited and Unsolicited Projects over Time

Expensive awards might be linked to agency solicitations. Figure 8, panel A, shows that before 2010 unsolicited RPGs had a central tendency towards greater costs,but since then solicited awards were more costly. Figure 8, panel B, shows that the proportion of solicited *projects* increased from 20 percent to 30 percent from FY1998 to FY2005, then remained stable until FY2016, and increased to 40 percent from 2016 to 2021. Meanwhile the proportion of *funds* going to solicited projects has steadily increased from 20 percent in FY1998 to 50 percent in FY2021.

**Figure 8:**
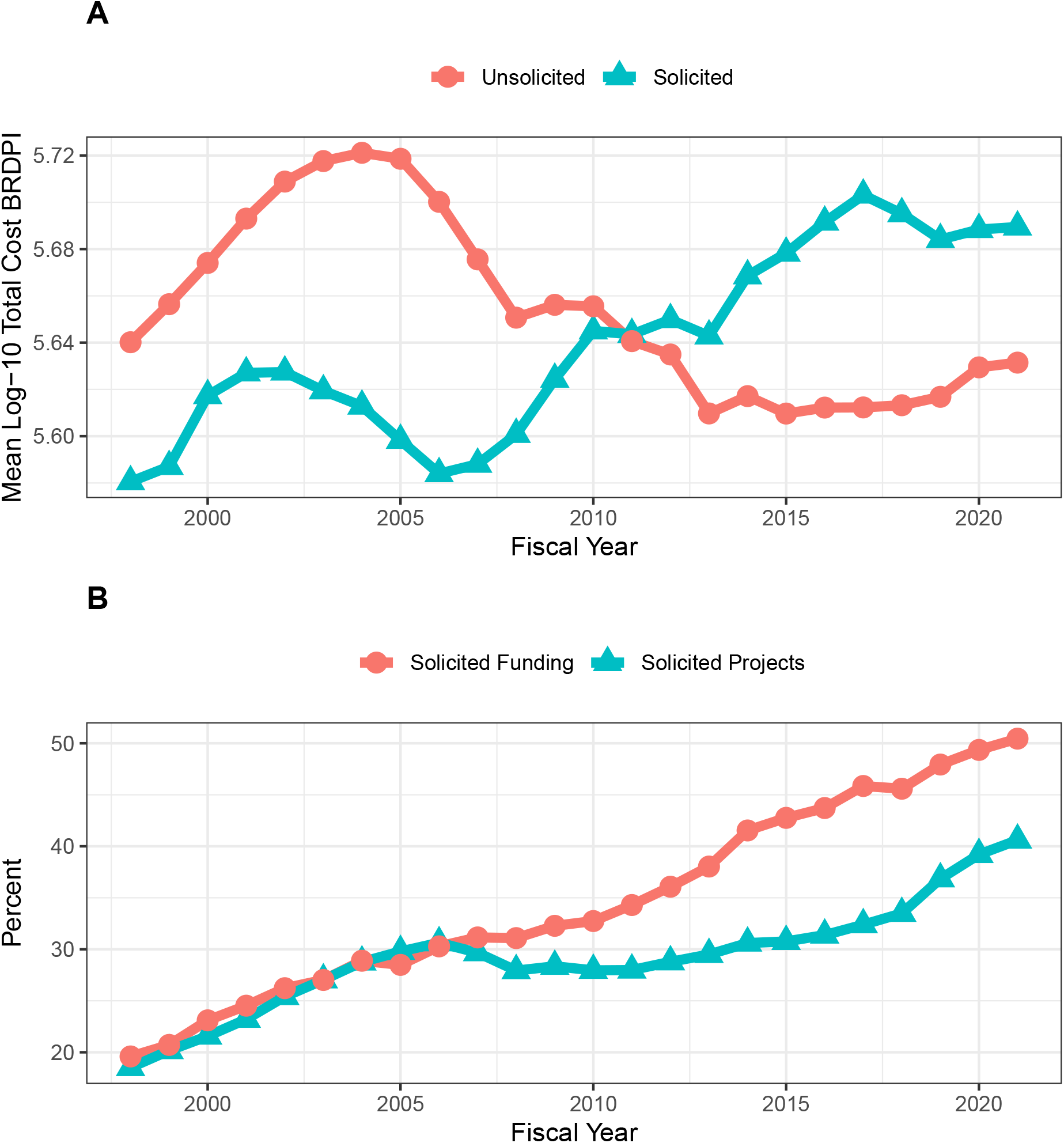
A: Log-transformed costs according to NIH solicitation. All costs are FY2021 BRDPI-adjusted. B: Percent of RPG projects and and percent of RPG funding going to solicited awards

Figure 9 shows box-plot distributions over time of log-transformed costs of unsolicited (panel A) and solicited (panel B) projects. Both groups of projects show variations in whisker ranges, but throughout time solicited projects have much greater degrees of variation (as reflected in larger whisker ranges). Table 2 compares solicited and unsolicited projects in FY2021 and FY2010. In FY2021 solicited projects were more expensive (mean of $710,000 versus $480,000), and more likely to be over $5 million, to be a cooperative agreement, to be a clinical trial, and to involve human participants. Solicited projects were also more likely to be funded through small R21 or R03 mechanisms, while much less likely to be funded via an R01-equivalent mechanism. Thus, the wide whisker ranges of solicited projects (Figure 9, panel B), which have become more common over time (Figure 8, panel B), may reflect both expensive and inexpensive awards. Inexpensive R21 and R03 awards have increased from 5 percent of projects in FY1998 to nearly 16 percent in FY2015, with a modest decline since (Figure 10).

**Figure 9:**
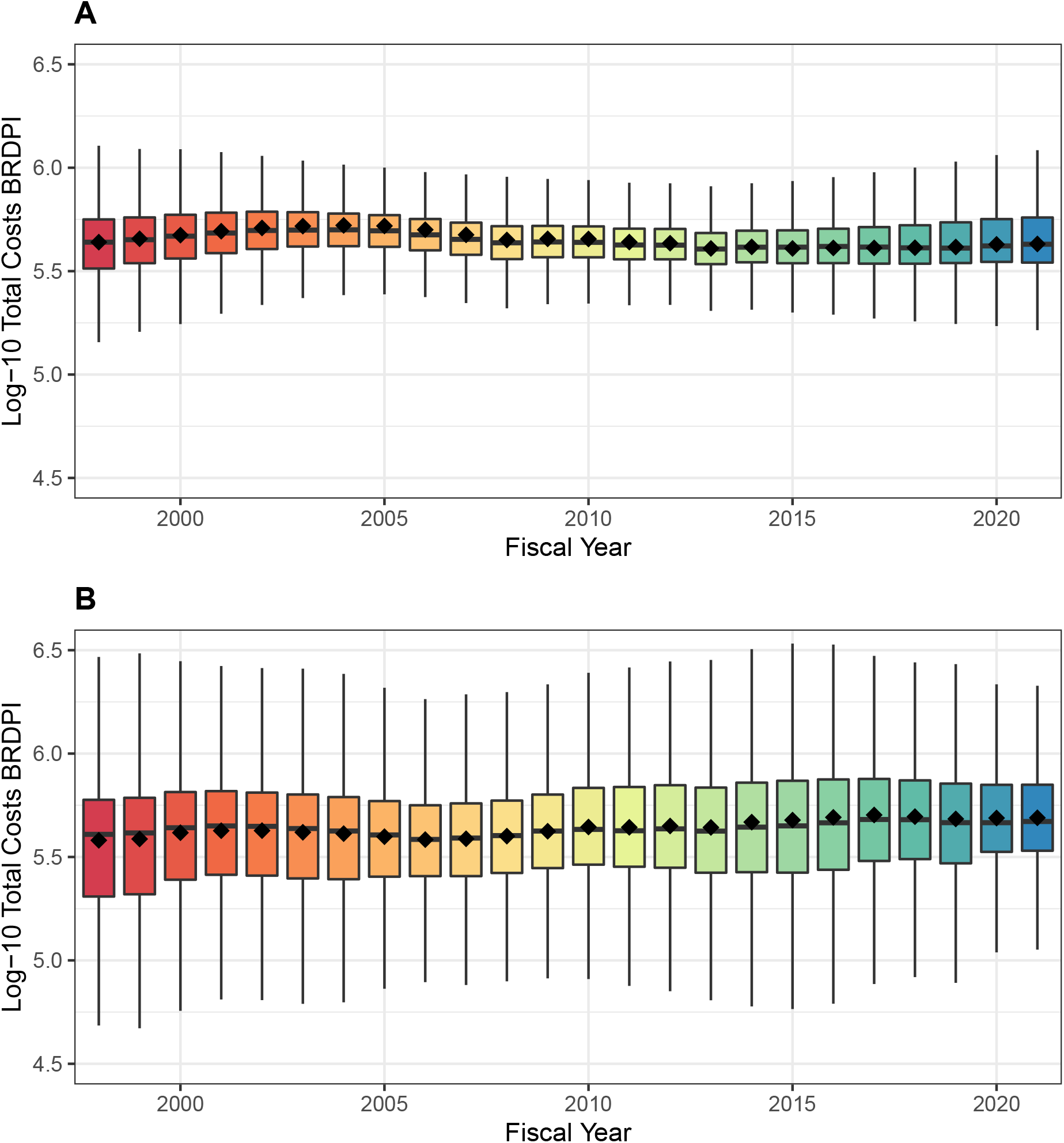
Box plot distributions of log-transformed costs for unsolicited (panel A) and solicited (panel B) NIH-funded RPGs, FY1998 to FY2021. The diamonds refer to mean values. Y-axes are similarly scaled for easier comparison. All costs are FY2021 BRDPI-adjusted.

**Figure 10:**
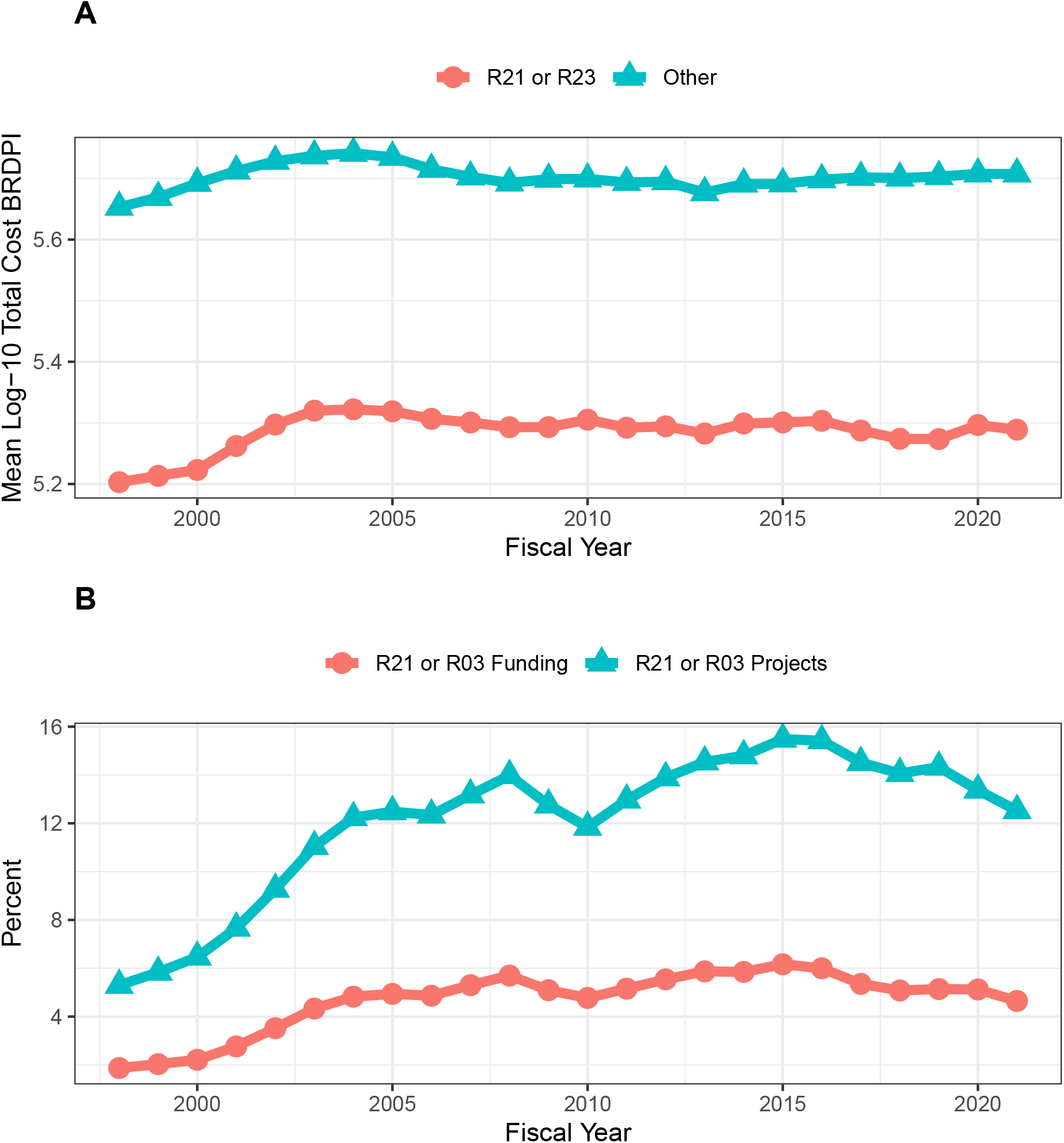
A: Costs according to R21 and R03 projects versus of all others. All costs are FY2021 BRDPI-adjusted. B: Percent of RPG projects and percent of RPG funding going to R21 and R03 projects

**Table 2:**
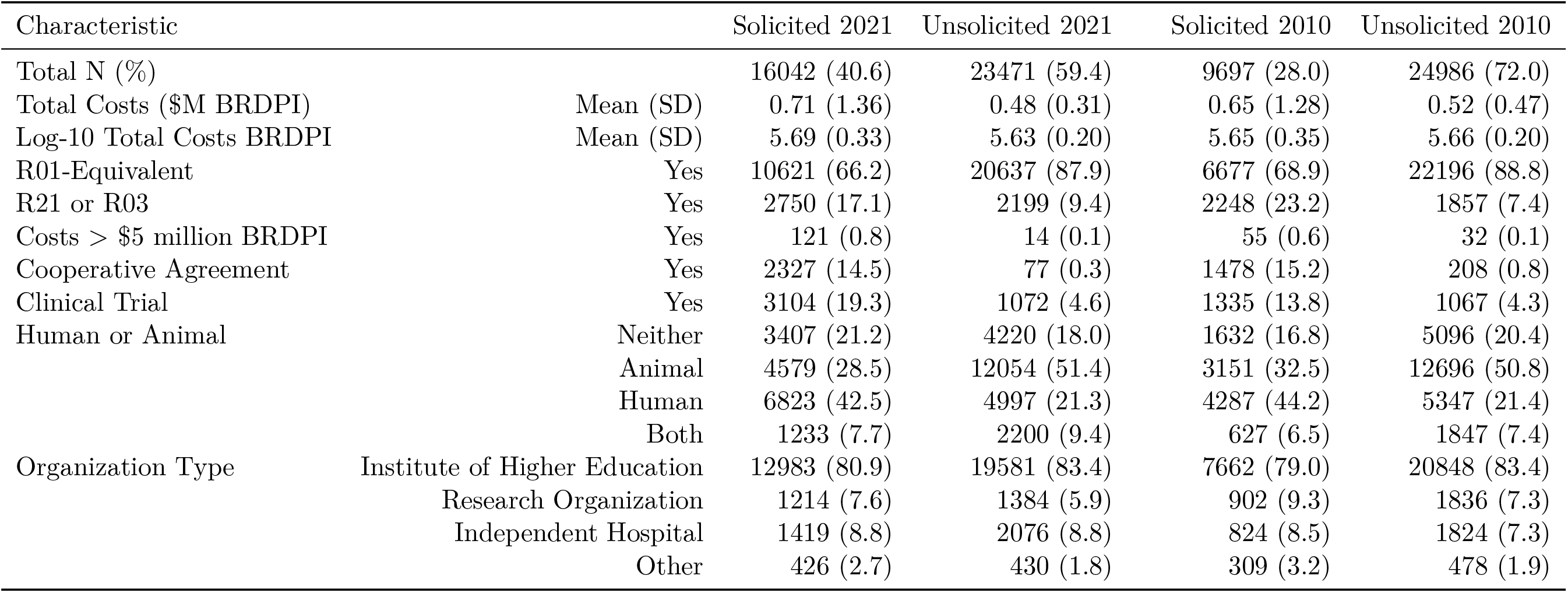
Characteristics of FY2021 and FY2010 RPGs according to solicitation

| Characteristic |  | Solicited 2021 | Unsolicited 2021 | Solicited 2010 | Unsolicited 2010 |
| --- | --- | --- | --- | --- | --- |
| Total N (%) |  | 16042 (40.6) | 23471 (59.4) | 9697 (28.0) | 24986 (72.0) |
| Total Costs (\$M BRDPI) | Mean (SD) | 0.71 (1.36) | 0.48 (0.31) | 0.65 (1.28) | 0.52 (0.47) |
| Log-10 Total Costs BRDPI | Mean (SD) | 5.69 (0.33) | 5.63 (0.20) | 5.65 (0.35) | 5.66 (0.20) |
| R01-Equivalent | Yes | 10621 (66.2) | 20637 (87.9) | 6677 (68.9) | 22196 (88.8) |
| R21 or R03 | Yes | 2750 (17.1) | 2199 (9.4) | 2248 (23.2) | 1857 (7.4) |
| Costs > \$5 million BRDPI | Yes | 121 (0.8) | 14 (0.1) | 55 (0.6) | 32 (0.1) |
| Cooperative Agreement | Yes | 2327 (14.5) | 77 (0.3) | 1478 (15.2) | 208 (0.8) |
| Clinical Trial | Yes | 3104 (19.3) | 1072 (4.6) | 1335 (13.8) | 1067 (4.3) |
| Human or Animal | Neither | 3407 (21.2) | 4220 (18.0) | 1632 (16.8) | 5096 (20.4) |
|  | Animal | 4579 (28.5) | 12054 (51.4) | 3151 (32.5) | 12696 (50.8) |
|  | Human | 6823 (42.5) | 4997 (21.3) | 4287 (44.2) | 5347 (21.4) |
|  | Both | 1233 (7.7) | 2200 (9.4) | 627 (6.5) | 1847 (7.4) |
| Organization Type | Institute of Higher Education | 12983 (80.9) | 19581 (83.4) | 7662 (79.0) | 20848 (83.4) |
|  | Research Organization | 1214 (7.6) | 1384 (5.9) | 902 (9.3) | 1836 (7.3) |
|  | Independent Hospital | 1419 (8.8) | 2076 (8.8) | 824 (8.5) | 1824 (7.3) |
|  | Other | 426 (2.7) | 430 (1.8) | 309 (3.2) | 478 (1.9) |

### Other RPG Characteristics and Costs over Time

Figure 11 shows that, as expected, RPGs involving clinical trials are more expensive (panel A) but, at least, over the last 10 years real costs remain stable. We acknowledge, though, that these analyses do not consider trial productivity measures, like numbers of patients enrolled. Figure 12 shows similar real-cost trends among RPGs irrespective of human or animal classification, though as expected projects involving human participants or human participants and animal models were more expensive than others.

**Figure 11:**
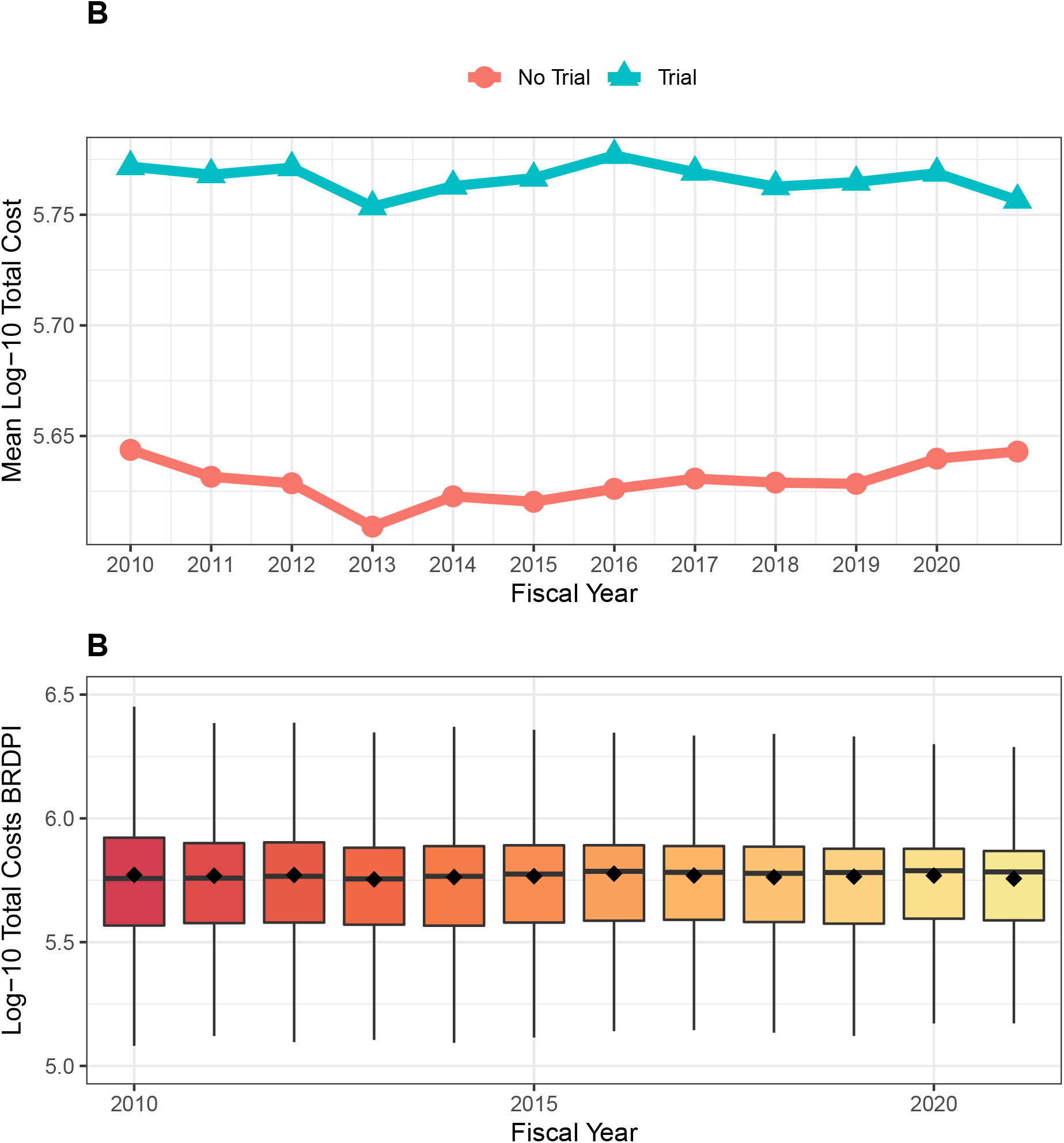
A: Log-transformed costs according to clinical trials for NIH-funded RPGs, FY2010 to FY2021. B: Box-plot distributions of log-transformed total costs for NIH-funded RPGs supporting clinical trials, FY2010 to FY2021. All costs are FY2021 BRDPI-adjusted.

**Figure 12:**
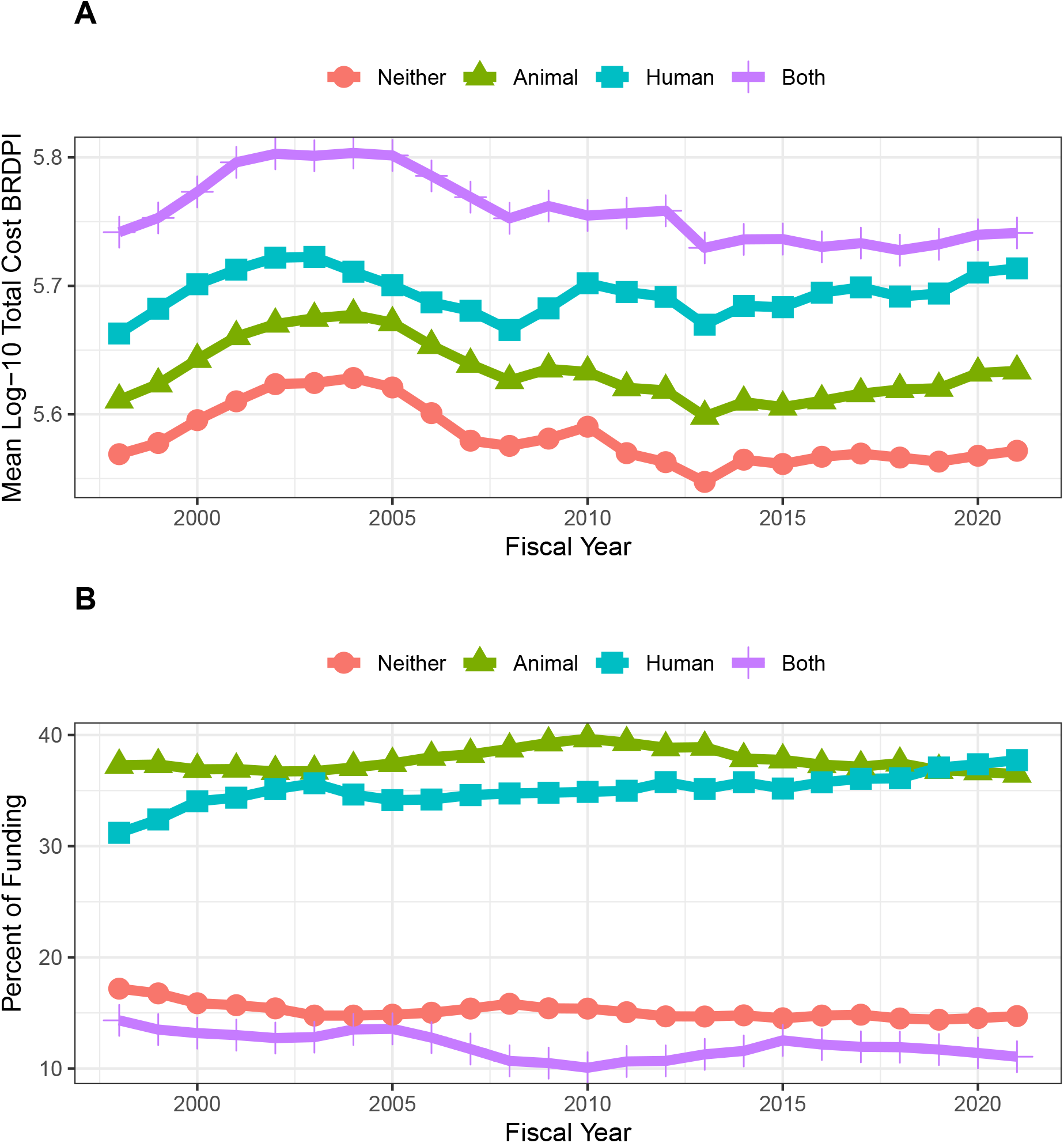
Panel A: Log-transformed costs of NIH-funded RPGs according to involvement of human participants and/or animal models, FY1998 to FY2021. All costs are FY2021 BRDPI-adjusted. Panel B: Proportion of RPG-funding going to different types of projects according to involvement of human participants and/or animal models, FY1998 to FY2021.

### Independent Association of Time with BRDPI-Adjusted RPG Costs

We conducted a series of regression analyses to examine whether there may be an association of time (that is fiscal year) with BRDPI-adjusted costs of RPG projects separate from those associated with funding mechanism, solicitation (or not), involvement of human participant or animal models, or type of recipient organization. We attempted multivariable linear regressions with log-transformed costs as the dependent variable, but upon inspection of residual diagnostics found poor model fit due to the fat-tailed distribution. We looked into other possible transformations (e.g. arcsinh, Box-Cox, center and scale, exponential, square-root, and Yeo-Johnson) and did not find substantive improvements. We therefore performed a wholly non-parametric random forest regression(Ishwaran and Kogalur, 2022) of log-transformed total costs; the model performed well, able to explain over 45 percent of the variance of costs. Time (that is fiscal year) did not contribute to prediction; that is, the variable importance of fiscal year was essentially zero. Figure 13 overlays the multivariable adjusted per-project cost with actual observed median costs and shows no material difference.

**Figure 13:**
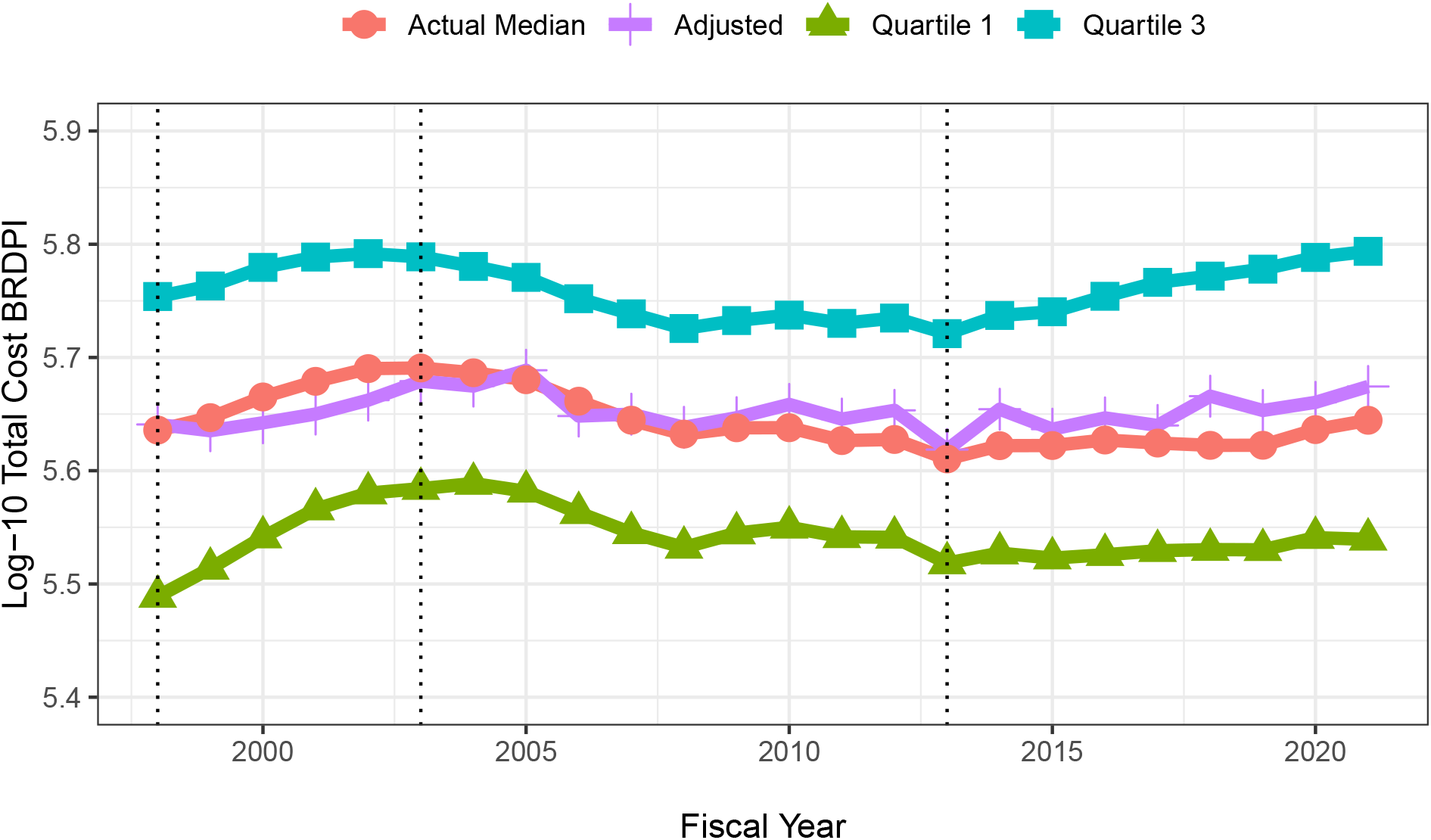
Actual and random-forest regression log-transformed costs for NIH-funded RPGs, FY1998 to FY2021. All costs are FY2021 BRDPI-adjusted.

## Discussion

The rate of inflation for NIH-funded research (that is the BRDPI) was higher than the general rate of inflation from 1998 until 2012; since then, the rate of inflation for NIH-funded research has been similar to the general rate of inflation. The more recent declines in the BRDPI may be related to caps on compensation of extramural investigators. Despite increases in nominal costs, recent years have seen increases in the absolute numbers of RPG and R01 awards. Real (BRDPI-adjusted) average and median RPG costs increased during the NIH-doubling (1998 to 2003), but have remained relatively stable since. Of note, though, the degree of variation of RPG costs has changed over time, with more marked extremes observed on both higher and lower levels of cost. On the higher end, over time NIH has been funding more cooperative agreements, more projects exceeding $5 million (in FY2021 BRDPI, *not nominal*, values), and more clinical trials. The top centile of projects are receiving a substantially greater share of the overall RPG funding pool. On the lower end of cost, over time the agency has been funding more low-cost mechanism awards (R03 and R21). On both ends of the cost spectrum, the agency is funding a greater proportion of solicited projects, with nearly half of RPG money going towards solicited projects. After adjusting for potential confounders in a wholly non-parametric machine learning regression, we find no independent association of time with BRDPI-adjusted costs. Recalling the automobile rental firm analogy, NIH may be pursuing the strategy of simultaneously purchasing more expensive (large vans) and less expensive (compact cars) vehicles, reflective of changing priorities and compositional effects over time.

### Why are Costs for Services (and Research) so High?

Increases in costs for research may be greater than increases in general economy-wide costs just as educational and health-care costs have increased at rates much greater than other costs. The Nobel-prize winning economist William Baumol explored differential increases in costs in his work on “the cost disease.”(Baumol and De Ferranti, 2012) The fundamental problem is that different sectors of the economy realize different rates of improvements in productivity. Baumol cites 4 musicians who play a Beethoven string quartet; there has been no change in productivity between 1826 and now. It takes just as many musicians just as much time to “produce” a live performance of a Beethoven string quartet. However, in other segments of the economy, productivity has increased dramatically, leading to increased wages for non-string-quartet workers. If we still want live performances of string quartets we have to pay much more now than in 1826 even though the output is unchanged because otherwise the musicians will choose other lines of work that pay more. The economists Eric Helland (Claremont McKenna College, RAND) and Alex Tabarrok (George Mason University) posted a report entitled “Why are the Prices so D-mn High?” in which they explain how Baumol’s construct works for explaining cost increases in the service sector, and in education and healthcare in particular.(Helland and Tabarrok, 2019)

Helland and Tabarrok illustrate the problem(Helland and Tabarrok, 2019)by imagining a simple two-product economy that produces only one good – cars – and one service – education. If society wants more education, the the opportunity cost (or price) will be fewer cars. Over time, productivity improves for both cars and education, but to a much greater degree for cars. If society wants to maintain the same ratio of education to cars, the price for that education relative to cars will be much higher. If society wants more education the price for education will be higher still. Thus, over time, relative prices for services (education) increase while prices for goods (cars) decline.

Bureau of Economic Analysis data(Helland and Tabarrok, 2019) on the relative costs of goods and services in the United States since 1950 show that the United States economy has shifted from goods to services while the relative prices for services (like education and healthcare) have increased. There is literature on the costs and productivity of research showing similar long-term patterns. For example, Scannell et al described “Eroom’s Law” of *declining efficiency* of pharmaceutical research and development dating back to 1950 and continuing relentlessly since.(Scannell et al.,2012) The number of drugs developed per billion dollars of R&D spending has *declined* by at least an order of magnitude. Other recent work has focused on the increasing costs of conducting clinical trials,(Sertkaya et al., 2016) whether sponsored by industry or by NIH.(Lauer et al., 2017) This literature identifies other drivers specific to pharmaceutical research or clinical trials in general, but these drivers may reflect general longstanding and inherent increases in the prices of services.

### Limitations

While we are able to describe changes in RPG costs over time, we note a number of important limitations. There is not a simple one-to-one link between specific grants and projects. Some projects are supported by multiple sources, including some outside of NIH. Individual grants are only partially able to cover costs, especially indirect costs for which recovery is nearly always partial. Because of salary caps and heterogeneous practices by which institutions use NIH funds for salary support, we do not have comprehensive information on compensation for personnel. Our regression analyses could only account for those variables we have in hand; nonetheless, the random forest model was able to account for a substantial proportion of the variance in RPG costs.

### Data and Materials

BRDPI and GDP-index values were obtained from the NIH Office of the Budget(NIH, 2022c). We queried Research Project Grant (RPG) data from NIH IMPAC II files. RPGs were defined as those grants with activity codes of DP1, DP2, DP3, DP4, DP5, P01, PN1, PM1, R00, R01, R03, R15, R21, R22, R23, R29, R33, R34, R35, R36, R37, R61, R50, R55, R56, RC1, RC2, RC3, RC4, RF1, RL1, RL2, RL9, RM1, SI2, UA5, UC1, UC2, UC3, UC4, UC7, UF1, UG3, UH2, UH3, UH5, UM1, UM2, U01, U19, U34 and U3R. Not all of these activity codes were used by NIH every year. For FY 2009 and 2010 we excluded awards made under the American Recovery and Reinvestment Act of 2009 (ARRA) and all ARRA solicited applications and awards. For FY2020 we excluded awards issued using supplemental Coronavirus (COVID-19) appropriations.

